# Duox generated reactive oxygen species activate ATR/Chk1 to induce G2 arrest in Drosophila tracheoblasts

**DOI:** 10.1101/2021.03.24.436759

**Authors:** Amrutha Kizhedathu, Piyush Chhajed, Lahari Yeramala, Deblina Sain Basu, Tina Mukherjee, Kutti R. Vinothkumar, Arjun Guha

## Abstract

Progenitors of the thoracic tracheal system of adult Drosophila (tracheoblasts) arrest in G2 during larval life and rekindle a mitotic program subsequently. G2 arrest is dependent on ATR-dependent phosphorylation of Chk1 that is actuated in the absence of detectable DNA damage. We are interested in the mechanisms that activate ATR/Chk1 (Kizhedathu et al., 2018, 2020). Here we report that levels of reactive oxygen species (ROS) are high in arrested tracheoblasts and decrease upon mitotic re-entry. High ROS is dependent on expression of Duox, an H_2_O_2_ generating-Dual Oxidase. ROS quenching by overexpression of Superoxide Dismutase 1, or by knockdown of Duox, abolishes Chk1 phosphorylation and results in precocious proliferation. Tracheae deficient in Duox, or deficient in both Duox and regulators of DNA damage-dependent ATR/Chk1 activation (Claspin/ATRIP/TOPBP1), can induce phosphorylation of Chk1 in response to micromolar concentrations of H_2_O_2_ in minutes. The findings presented reveal that H_2_O_2_ activates ATR/Chk1 in tracheoblasts by a non-canonical, potentially direct, mechanism.

## INTRODUCTION

Ataxia Telangiectasia Mutated-Related Kinase (ATR, *Mei-41*) and its substrate, Checkpoint Kinase 1 (Chk1, *Grapes*), are essential for DNA damage repair and for normal development (Artus & Cohen-Tannoudji, 2008; Blythe & Wieschaus, 2015; Cimprich & Cortez, 2008). The mechanism for activation of the ATR-Chk1 axis in response to DNA damage, and the mechanism by which activated Chk1 induces cell cycle arrest, have been well characterized (Choi et al., 2010; Cimprich & Cortez, 2008; Delacroix et al., 2007; Lee et al., 2012; Xu & Leffak, 2010). In contrast, the mechanisms for the activation of ATR/Chk1 during development, and the roles of these proteins therein, are less well understood. We reported recently that the ATR/Chk1 axis is co-opted during Drosophila development for inducing G2 arrest in progenitor cells (Kizhedathu et al., 2018, 2020). The current study was designed to shed light on the mechanism of activation of the ATR-Chk1 axis in this context, and to probe if it is distinct from the mechanism for DNA damage-induced activation.

The progenitors of the adult tracheal (respiratory) system in Drosophila remain quiescent through larval life and rekindle a mitotic program at the onset of pupariation. Progenitors of the tracheal branches of the second thoracic metamere in the adult (Tr2, hereafter referred to tracheoblasts) are arrested in the G2 phase of the cell cycle for ∼56 h and enter mitoses thereafter. Pertinently, G2 arrest in tracheoblasts is dependent on ATR-dependent phosphorylation of Chk1. Two aspects of the mechanism of G2 arrest are striking and merit mention here. First, arrested tracheoblasts phosphorylate Chk1 (phosphorylated Chk1, pChk1) in the absence of detectable DNA damage. Second, arrested tracheoblasts upregulate Chk1 expression in a Wnt signaling-dependent manner and high levels of Chk1 expression are essential for arrest (Kizhedathu et al., 2018, 2020). How the ATR/Chk1 pathway is activated in tracheoblasts, in the absence of DNA damage, was unclear from these studies.

ATR together with ATM (Ataxia Telangiectasia Mutated) and DNA-PK are the principal sensors of DNA damage and effectors of the DNA damage response (Durocher & Jackson, 2001). Interestingly, there is evidence that these kinases can also be activated by non-canonical mechanisms that are not dependent on DNA damage (Guo et al., 2010). Reactive Oxygen Species (ROS) are a group of oxygen-derived small molecules that interact avidly with macromolecules like proteins, lipids and nucleic acids and alter their function (Corcoran & Cotter, 2013b). Importantly, there is evidence that ROS can modulate the activity of the aforementioned kinases by both canonical and non-canonical mechanisms. High levels of ROS in cells can cause DNA damage. In addition, ROS has been shown to directly activate ATM by stabilizing ATM homodimers through the formation intermolecular disulphide bridges (Guo et al., 2010). ROS-activated ATM can phosphorylate and activate substrates like Checkpoint Kinase 2 (Chk2). The study on ATM alerted us to the possibility that there are non-canonical modes of kinase activation and ROS-based activation is one such mechanism.

ROS have also emerged as important regulators of developmental processes (Zhang et al., 2016). The regulation of ROS levels in cells during development is orchestrated by altering rates of aerobic respiration in cells or by altering the levels of expression of ROS-producing enzymes like NADPH oxidases (NOXs). Interestingly, the Drosophila genome encodes two NADPH oxidases - NOX and Dual oxidase (Duox) (Kim & Lee, 2014) and one of these, Duox, is expressed at high levels in the larval tracheal system (*Project: FlyAtlas-RNA.Larva*, n.d.). Duox catalyzes the oxidation of NADPH leading to the generation of H_2_O_2_ (Bedard & Krause, 2007; Geiszt et al., 2003).

Our interest in the mechanism for activation of ATR/Chk1 led us to probe the levels of ROS in tracheoblasts during larval life. We found the levels were high when the cells were arrested and low after the cells rekindled cell division. We quenched ROS levels in tracheoblasts via overexpression of the antioxidant Superoxide Dismutase 1 to find that the cells started dividing precociously. These findings led us in turn to probe the role of Duox in the regulation of ROS and the mechanism by which ROS regulate the ATR/Chk1 axis.

## RESULTS

### High ROS is required for G2 arrest in larval tracheoblasts

The cells that comprise the tracheal branches of the second thoracic metamere (Tr2) of the larvae are differentiated adult progenitors that contribute to the development of adult tracheal structures (Guha et al., 2008; Guha & Kornberg, 2005). The cells that make up the Dorsal Trunk (DT) in Tr2, hereafter referred to as tracheoblasts, are the focus of our studies. Tracheoblasts remain arrested in the G2 phase of the cell cycle during larval life and initiate mitosis thereafter. Analysis of cell cycle phasing of tracheoblasts using FUCCI has shown that the cells are in the G1 phase at the time the embryo hatches into a larva and that the cells transition from G1-S-G2 in the first larval instar (L1). Tracheoblasts remain in G2 from the second larval instar (L2) through to mid third larval instar (L3) (32-40 h L3, ∼56 h) and divide rapidly thereafter ((Kizhedathu et al., 2018, 2020), Figure 1 A).

**Figure 1:**
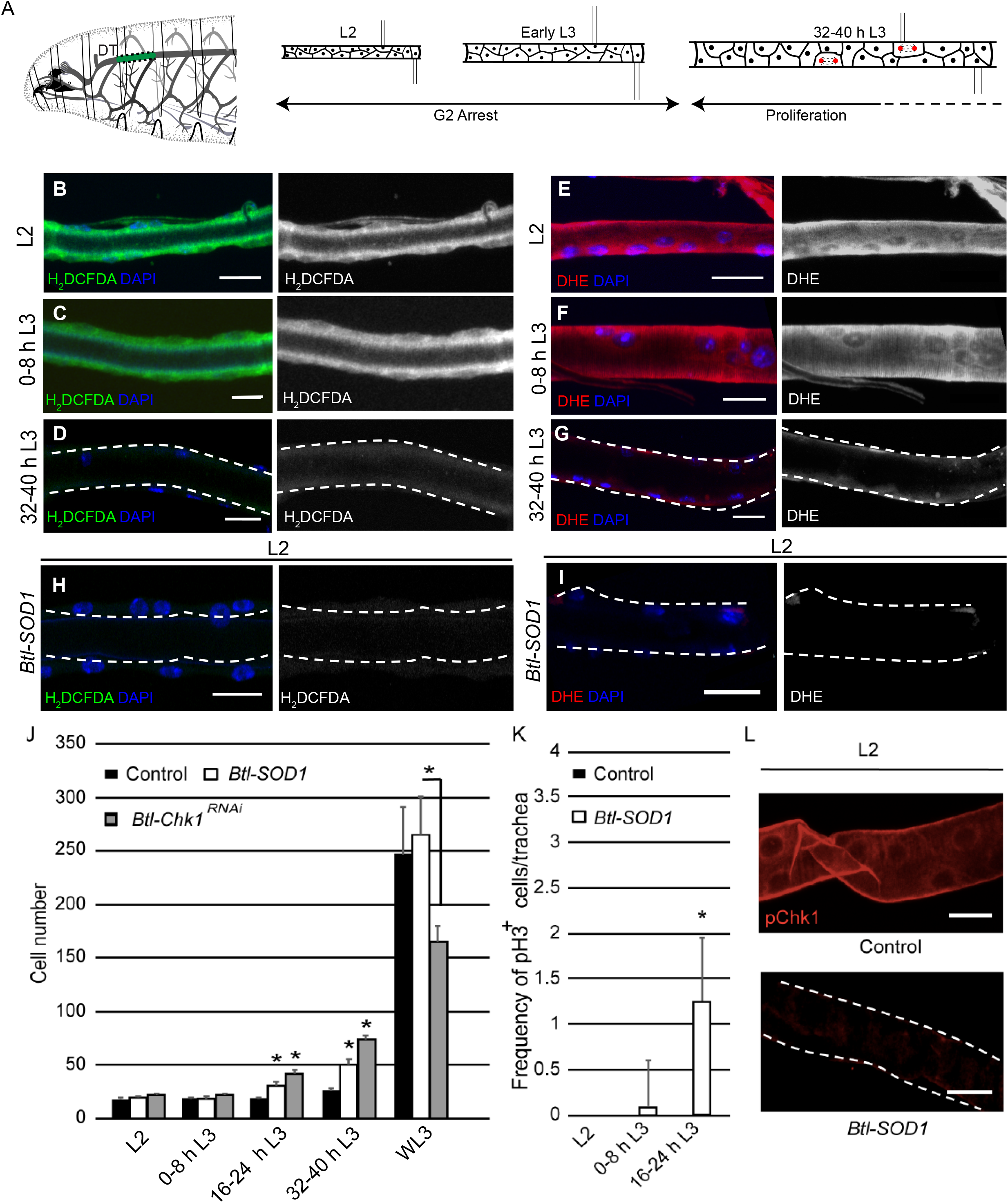
High levels of Reactive Oxygen Species (ROS) are required for Chk1 activation and G2 arrest in tracheoblasts. **(A)** A diagram of the third instar larva showing the dorsal trunk (DT) of the second thoracic metamere (Tr2) colored in green and marked by dashed line. The cartoon also shows the timecourse of G2 arrest and cell division in Tr2 tracheoblasts that comprise the DT of the larva. The cells in Tr2 DT remain geographically isolated from tracheal cells in other branches during larval life. **(B-D)** Levels of the ROS reporter 2’,7’-dichlorodihydrofluorescein diacetate (H_2_DCFDA) in Tr2 DT during larval stages. Shown in the figures are H_2_DCFDA staining in L2 **(B)**, 0-8 h L3 **(C)** and 32-40 h L3 **(D). (E-G)** Levels of the ROS reporter Dihydroethidium (DHE) in Tr2 DT during larval stages. Shown in the figures are DHE staining in L2 **(E)**, 0-8 h L3 **(F)** and 32-40 h L3 **(G)**. **(H-I)** Effect of *Btl*-Gal4-dependent overexpression of Superoxide Dismutase 1 (SOD1) on levels of ROS reporters in Tr2 DT. **(H)** H_2_DCFDA staining in *Btl-SOD1* (*btl-GAL4/UAS-SOD1)* expressing larvae (n ≥ 6 tracheae per condition per timepoint). **(I)** DHE staining in *Btl-SOD1* larvae (n ≥ 6 tracheae per condition per timepoint). **(J)** Effect of SOD1 overexpression on cell numbers in Tr2 DT at different larval stages. Graph shows numbers of Tr2 tracheoblasts in wild type *(btl-Gal4), Btl-SOD1 (btl-GAL4/UAS-SOD1)* and *Btl-Chk1^RNAi^ (btl-GAL4/UAS- Chk1^RNAi^)* larvae at L2, 0-8 h L3, 16-24 h L3, 32-40 h L3 and wandering L3 (WL3) (mean values ± standard deviation, n ≥ 7 tracheae per condition per timepoint). **(K)** Effect of SOD1 overexpression on mitotic indices in Tr2 DT (see text). Graph shows mitotic indices in Tr2 DT in wild type and *Btl-SOD1 (btl-GAL4/UAS-SOD1)* expressing larvae at L2, 0-8 h L3 and 16-24 h L3 (mean values ± standard deviation, n ≥ 7 tracheae per condition per timepoint). **(L)** Effect of SOD1 overexpression on Chk1 phosphorylation in Tr2 tracheoblasts. Shown in the figure is phosphorylated Chk1 (pChk1, phospho-Chk1Ser^345^) immunostaining (red) in Tr2 DT in wild type *(btl-GAL4)* and *Btl-SOD1 (btl-GAL4/UAS-SOD1)* larvae at L2. Scale bars = 10 µm. Student’s paired t-test: *p<0.05.

To probe the role of ROS in the regulation of G2 arrest in tracheoblasts, we assessed the levels of mitochondrial and cytoplasmic ROS in tracheoblasts at L2, early L3 and 32-40 h L3 using compartment-specific ROS sensors. Mitochondrial ROS levels were measured using a genetically encoded, mitochondrial targeted, redox sensitive GFP-fusion protein (MTS roGFP2) (Liu et al., 2012). Redox sensitive GFPs (roGFPs) are generated by substituting surface exposed amino acids with cysteines that can form disulfide bridges, making the protein sensitive to redox modifications. The formation of disulfide bridges results in blue-shifted light emission that can be readily quantified (Dooley et al., 2004; Hanson et al., 2004). Cytoplasmic ROS levels were measured using two well-established, redox sensitive dyes: 2′,7′-Dichlorodihydrofluorescein diacetate (H_2_DCFDA) and Dihydroethidium (DHE). Both H_2_DCFDA and DHE are cell permeable molecules that alter light emission upon oxidation (Yang et al., 2014).

To assay the levels of mitochondrial ROS, we expressed MTS roGFP2 in the trachea using the Breathless (*Btl*) promoter, dissected trachea at different timepoints and imaged the samples on a confocal microscope. Each sample was excited at 405 nm (oxidized GFP) and 488nm (reduced GFP) and ratios of the intensities of emission at 535 nm were calculated from images obtained (405/488 fluorescence ratio). The ratio was higher in L2, than at 0-8 h L3 or 32-40 h L3 (Figure1-Figure supplement 1, n ≥ 6 tracheae per condition per experiment, n = 3). We inferred that mitochondrial ROS levels are higher in L2 than at 0-8 h L3 or 32-40 h L3. Next, we probed the levels of cytoplasmic ROS. Analysis of H_2_DCFDA and DHE staining in tracheoblasts at various stages revealed that levels of both reporters are readily detectable at L2 (Figure 1 B, E, n ≥6 tracheae per condition per experiment, n = 3) and 0-8 h L3 (Figure 1 C, F, n ≥ 6 tracheae per condition per experiment, n = 3) and nearly undetectable at 32-40 h L3 (Figure 1 D, G, n ≥ 6 tracheae per condition per experiment, n = 3). Taken together, the analysis of ROS reporters showed that dynamics of mitochondrial and cytoplasmic ROS are different. Pertinently, these studies showed that dynamics of cytoplasmic ROS paralleled G2 arrest and mitotic re-entry.

Next we asked whether changes in ROS levels had any bearing on the cell cycle program. To answer this question, we characterized a genetic approach for quenching ROS in tracheoblasts. Superoxide dismutase 1 (SOD1) is a cytoplasmic enzyme that scavenges ROS (Blackney et al., 2014). Thus, we overexpressed SOD1 in trachea *(Btl-*SOD1) and examined the effects of SOD1 overexpression on ROS levels by H_2_DCFDA and DHE staining. Levels of H_2_DCFDA and DHE were found to be significantly lower in *Btl-SOD1* expressing animals compared to controls (Figure 1 H-I, compare with Figure 1 B, E, n ≥ 6 tracheae per condition per experiment, n = 3). This showed that SOD1 overexpression is an effective way to quench ROS in tracheae. We then determined whether SOD1 overexpression altered the cell cycle program of tracheoblasts. We counted the number of cells of tracheoblasts in Tr2 DT at L2, 0-8 h L3, 16-24 h L3, 32-40 h L3 and Wandering L3 (WL3) and quantified the frequencies of phospho-Histone H3^+^ (pH3^+^) mitotic figures in Tr2 DT at L2, 0-8 h L3 and 16-24 h L3. Analysis of the frequencies of cells and pH3^+^ figures showed that SOD1 overexpression resulted in precocious cell division from 0-8 h L3 (Figure 1 J, K, n ≥ 7 tracheae per timepoint, Figure 1 – Source data 1,2). We noted that the timecourse of cell divisions in *Btl-*SOD1 tracheae is similar to that of Chk1 mutant animals (*Btl-*Chk1^RNAi^) (Figure 1 J). Based on these data we concluded that quenching of ROS phenocopies the loss of Chk1 in these cells. We also observed a significant difference in tracheoblast behavior in *Btl-*Chk1^RNAi^ and *Btl-SOD1* expressing animals. Tracheoblasts in *Btl-*Chk1^RNAi^ rekindle mitoses earlier than wild type but divide more slowly thereafter. In contrast, tracheoblasts in *Btl-SOD1* rekindle mitoses earlier than wild type but do not appear to divide more slowly than wild type cells (see Discussion).

The findings above led us to investigate the levels of phosphorylated (activated) Chk1 in tracheoblasts in wild type and *Btl-*SOD1 animals. As reported previously, pChk1 levels are high in L2 and early L3 and diminished at 32-40 h L3. pChk1 immunostaining in *Btl-SOD1* expressing tracheae in L2 and early L3 showed that pChk1 levels were reduced in comparison to wild type at these respective stages (Figure 1 K, data not shown, n ≥ 6 tracheae per condition per experiment, n = 3). We inferred that high ROS levels are necessary for G2 arrest and that high ROS contributes in some manner to high levels of pChk1.

### High ROS in G2-arrested tracheoblasts is dependent on *Duox*

The identification of ROS as regulator of G2 arrest in tracheoblasts raises two questions. First, how are ROS levels regulated in the tracheoblasts? Second, how does high ROS translate into high levels of pChk1? As mentioned previously, the H_2_O_2_-generating enzyme Duox is expressed at high levels in larval tracheae (Bedard & Krause, 2007; Geiszt et al., 2003). To probe the role of Duox in the generation of ROS in tracheoblasts, we first examined Duox expression in tracheoblasts at different larval stages using quantitative RT-PCR (qPCR). We isolated mRNA from micro-dissected fragments of Tr2 DT at different timepoints and utilized these samples to query Duox expression. Our analysis showed that Duox mRNA levels are higher at L2 and 0-8 h L3 than at 32-40 h L3 (Figure 2 A, n ≥ 15 tracheal fragments per timepoint per experiment, n = 3 experiments). We concluded that the timecourse of Duox mRNA expression correlates with the timecourse of ROS accumulation in the tracheae.

**Figure 2:**
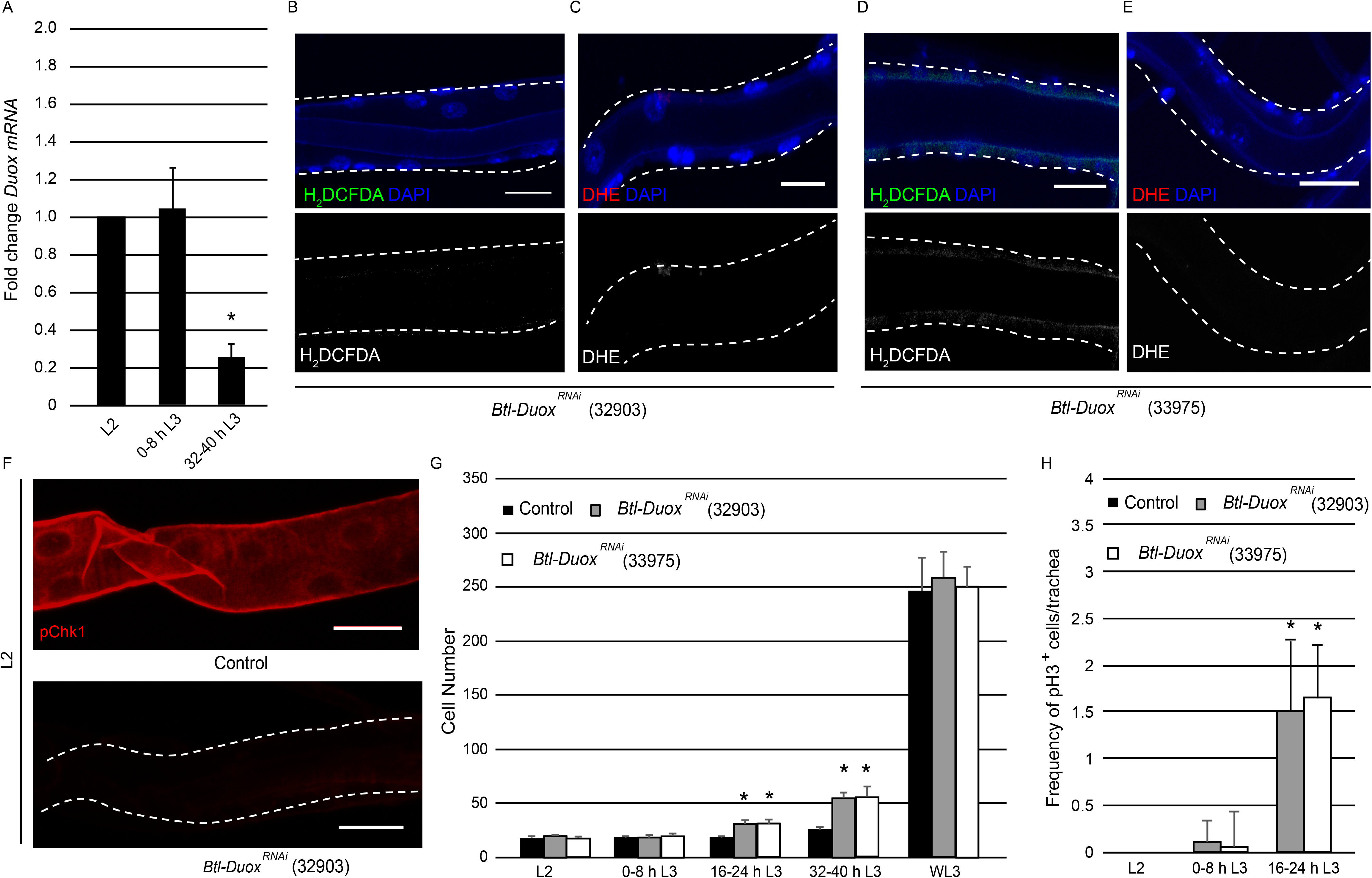
High ROS in tracheoblasts is dependent on *Duox* expression. **(A)** Quantitative PCR analysis of *Duox* mRNA levels in micro-dissected Tr2 DT fragments at different stages. Graph shows fold change in *Duox* mRNA in Tr2 DT fragments from wild type (*btl- GAL4*) larvae at L2, 0-8 h L3 and 32-40 h L3. Fold change has been represented with respect to L2 (n = 3 experiments, n ≥ 15 Tr2 DT fragments/stage/experiment, mean ± standard deviation). **(B-E)** Effect of the knockdown of *Duox* expression on the levels of ROS reporters in Tr2 DT. Shown here are the results of the expression of two different Duox RNAi lines (32903 and 33975). **(B, D)** H_2_DCFDA staining and **(C, E)** DHE staining in Tr2 DT in *Btl-Duox^RNAi^ (btl-GAL4/+; UAS-Duox^RNAi^(32903)/+* **(B,C)** and *btl- GAL4/+; UAS-Duox^RNAi^(33975)/+* **(D,E)**) larvae at L2 (n ≥ 6 tracheae per condition per timepoint) **(F)** Effect of reduction of *Duox* expression on levels of pChk1 in Tr2 DT. pChk1 immunostaining (red) in Tr2 DT in wild type *(btl-Gal4,* same as figure 1 L) and *Btl-Duox^RNAi^ (btl-GAL4/+; UAS- Duox^RNAi^(32903)/+)* larvae at L2 **(G)** Effect of the knockdown of *Duox* expression on cell numbers in Tr2 DT at different larval stages. Graph shows cell numbers of Tr2 tracheoblasts in wild type *(btl-Gal4,* Same as Figure 1 J) and *Btl-Duox^RNAi^ (btl- GAL4/+; UAS-Duox^RNAi^(32903)/+* and *btl-GAL4/+; UAS-Duox^RNAi^(33975)/+)* larvae at L2, 0- 8 h L3, 16-24 h L3, 32-40 h L3 and WL3 (mean values ± standard deviation, n ≥ 7 tracheae per condition per timepoint). **(H)** Effect of the knockdown of *Duox* expression on mitotic indices in Tr2 DT. Graph shows mitotic indices in Tr2 DT in wild type *(btl-Gal4)* and *Btl- Duox^RNAi^ (btl-GAL4/+; UAS-Duox^RNAi^(32903)/+* and *btl-GAL4/+; UAS-Duox^RNAi^(33975)/+)* larvae at L2, 0-8 h L3 and 16-24 h L3 (mean values ± standard deviation, n ≥ 7 tracheae per condition per timepoint). Scale bars = 10 µm. Student’s paired t-test: *p<0.05.

To probe whether Duox is the driver of ROS accumulation, we knocked down levels of Duox in the tracheal system by RNA interference and examined the levels of ROS reporters. The reduction in the levels of Duox using two different RNAi lines (BDSC-32903, BDSC-33975) followed by H_2_DCFDA and DHE staining showed that the knockdown of Duox leads to a dramatic decrease in levels of both reporters in comparison to wild type (Figure. 2 B-E, compare with Figure 1B and E, n ≥ 6 tracheae per condition per experiment, n = 3). Based on these data we inferred that the high levels of ROS in arrested tracheoblasts are dependent on Duox expression.

Next, we examined how the knockdown of Duox impacted Chk1 phosphorylation and the program of cell division. Consistent with the previous findings with SOD1 overexpression, we observed that the knockdown of Duox resulted in the loss of pChk1 (Figure 2 F, n ≥ 6 tracheae per condition per experiment, n = 3). We subsequently counted the number of cells of tracheoblasts at L2, 0-8 h L3, 16-24 h L3, 32-40 h L3 and Wandering L3 (WL3) and quantified the frequencies of pH3^+^ nuclei in Tr2 DT at L2, 0-8 h L3 and 16-24 h L3. We found that *Btl-Duox^RNAi^* expressing animals rekindle cell divisions sooner than their wild type counterparts. Interestingly, *Btl-Duox^RNAi^* expressing animals also showed no obvious slowdown in cell division rate after mitotic re-entry (Figure 2 G, H, n ≥ 7 tracheae per timepoint, Figure 2 – Source data 1,2).

### ROS dependence reveals a novel mode of ATR/Chk1 regulation

Having identified the source for high ROS in arrested tracheoblasts, we turned our attention to addressing how ROS was coordinating G2 arrest. Our previous studies have shown that Wnt-dependent transcriptional upregulation of Chk1 is essential for G2 arrest. Wnt signaling in the trachea is mediated by four Wnt ligands: Wg, Wnt5, Wnt6 and Wnt10 that are expressed by the tracheoblasts (Figure 3A). All ligands are expressed at high levels in arrested cells but downregulated post mitotic entry. We have also shown that the four Wnts act synergistically to upregulate Chk1 expression but are redundant for expression of other Wnt targets like Fz3.

**Figure 3:**
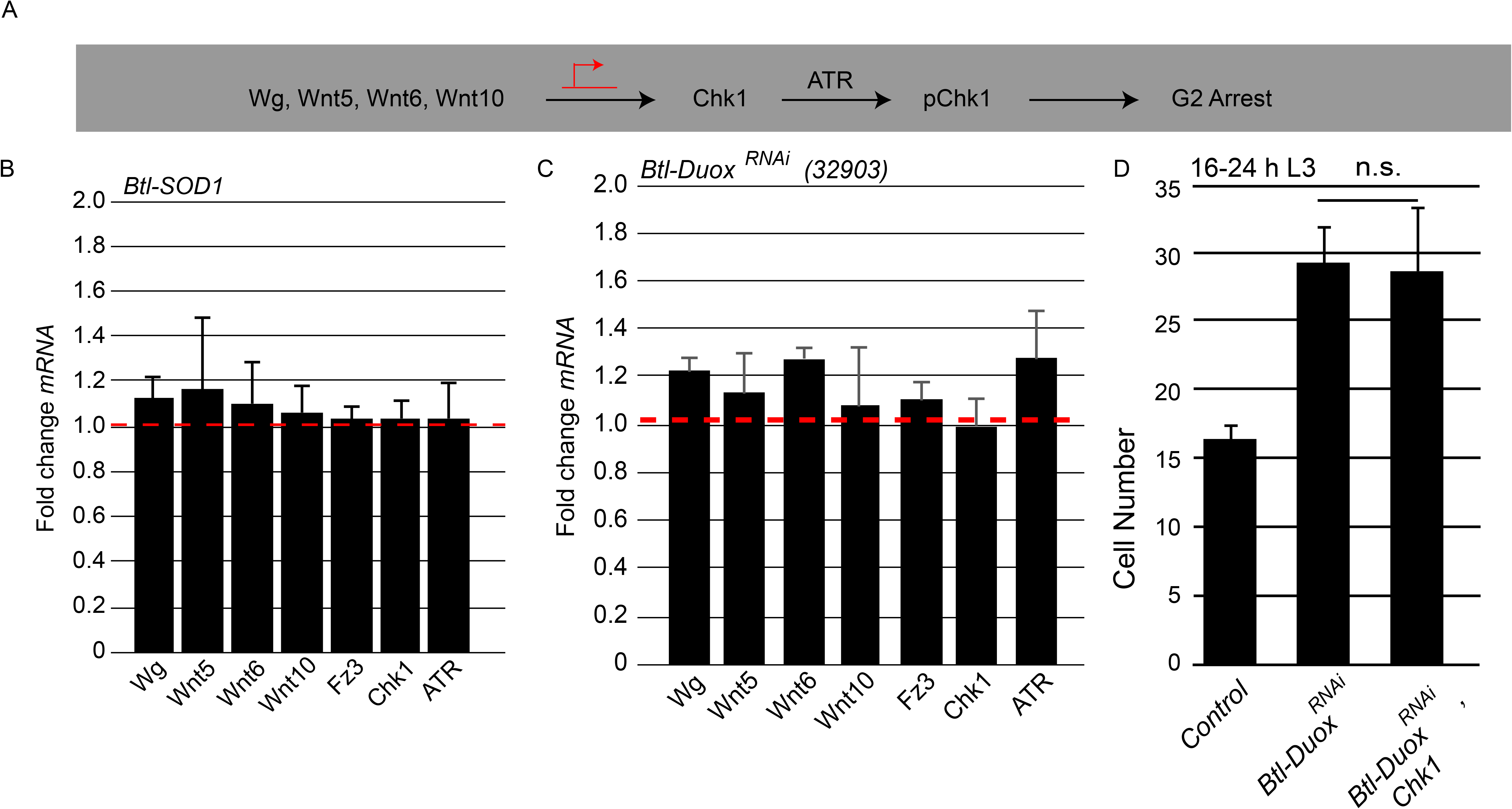
ROS dependence identifies a novel pathway for the regulation of ATR/Chk1 in tracheoblasts. **(A)** Model for G2 arrest mechanism in Tr2 tracheoblasts based on previous studies. Earlier work has shown that four Wnt ligands (*Wg, Wnt5, Wnt6, Wnt10*) act synergistically to upregulate Chk1 mRNA levels in arrested tracheoblasts. High levels of Chk1 expression are necessary for G2 arrest and Chk1 overexpression can rescue defects in Wnt signaling (Kizhedathu et al., 2020). **(B-C)** Effect of SOD1 overexpression and Duox knockdown on expression of Wnts and Wnt-target genes. Quantitative PCR analysis of levels of *Wg, Wnt5, Wnt6, Wnt10, Fz3, Chk1 and ATR* mRNA in micro-dissected Tr2 DT fragments at L2. Graph shows fold change in *Wg, Wnt5, Wnt6, Wnt10, Fz3, Chk1 and ATR* mRNA levels in Tr2 DT fragments expressing *Btl-SOD1 (btl-GAL4/UAS-SOD1)* **(B)** and *Btl- Duox^RNAi^ (btl-GAL4/+; UAS-Duox^RNAi^(32903)/+)* **(C)**. Fold change has been represented with respect to wild type *(btl-Gal4)* represented by dashed red line at L2 (n = 3 experiments, n ≥ 15 Tr2 DT fragments/stage/experiment, mean ± standard deviation). **(D)** Effect of Chk1 overexpression on cell numbers in Tr2 DT in *Btl-Duox^RNAi^* larvae at 16-24 h L3. Graph shows numbers of Tr2 tracheoblasts in wild type *(btl-Gal4)*, *Btl-Duox^RNAi^* (*btl-GAL4/+; UAS- Duox^RNAi^(32903)/+)* and *Btl-Duox^RNAi^, Chk1* (*btl-GAL4/+; UAS-Duox^RNA^(32903)^i^/ UAS-Chk1)* larvae at 16-24 h L3 (mean values ± standard deviation, n ≥ 7 tracheae per condition per timepoint). Student’s paired t-test: n.s (not significant).

Our first step toward characterizing the role of ROS in G2 arrest was to analyse how ROS levels impacted Wnt signaling and the expression of Wnt target genes, particularly Chk1. We micro-dissected Tr2 fragments from *Btl-SOD1* and *Btl-Duox^RNAi^* expressing animals at L2, extracted mRNA and analyzed expression of both Wnts and Wnt target genes by qPCR. We found that the expression of all Wnt ligands, Fz3 and Chk1 were comparable in wild type, *Btl-SOD1* and *Btl-Duox^RNAi^* expressing animals (Figure 3 B-C, Red dashed line marks wild type levels, n ≥ 15 tracheal fragments per timepoint per experiment, n = 3 experiments). This showed that perturbations in ROS levels in the trachea do not impact Wnt signaling nor expression of Wnt targets like Chk1. These findings implied that ROS must regulate activation of the ATR/Chk1 by some other mechanism. We also assayed the levels of ATR in *Btl-SOD1* and *Btl-Duox^RNAi^* expressing animals by qPCR and found no change in ATR transcript levels compared to control (Figure 3 B-C, Red dashed line marks wild type levels). Together, the qPCR data suggested that ROS did not regulate the abundance either ATR or Chk1 transcripts.

We have shown previously that precocious mitotic re-entry observed in Wnt signaling-deficient tracheoblasts (*Btl-TCF^RNAi^*) can be rescued by overexpression of Chk1 (Kizhedathu et al., 2020). Thus, to further confirm the ROS levels do not regulate levels Chk1 levels, we overexpressed Chk1 in tracheoblasts (Btl-Chk1) and examined cell proliferation in *Btl-Duox^RNAi^* expressing animals. We counted numbers of tracheoblasts in Btl*-Duox^RNAi^, Btl-Chk1* animals at 16-24 h L3 to find that the numbers were considerably higher than wild type and comparable to the numbers in *Btl-Duox^RNAi^* animals (Figure 3 D, n ≥ 7 tracheae per timepoint, Figure 3 – Source data 1). Thus, the data suggested that ROS does not activate ATR/Chk1 by facilitating Wnt signaling and Chk1 overexpression.

### ROS-dependent activation of ATR/Chk1 does not require ATRIP/TOPBP1/Claspin

The coincidence of high ROS levels and activated Chk1 in cells would typically suggest that ROS-dependent genotoxic stress was leading to activation of Chk1. However, our analysis of DNA damage in tracheoblasts, using the well characterized marker for double strand DNA breaks (γ-H2AX), has shown that there is no detectable DNA damage in arrested cells (Kizhedathu et al., 2018). In light of the findings with respect to ROS, we decided to probe more rigorously the incidence of genotoxic stress in tracheoblasts and the role of the DNA damage response in Chk1 activation.

Next we re-examined the levels of DNA damage in tracheoblasts using two assays. First, we examined the accumulation of 8-Oxo-2’-deoxyguanosine (8-oxo-dG), a marker for nucleotide oxidation. Second, we scored the frequencies of nuclear foci of RPA70, a protein that binds single strand DNA breaks. To validate nuclear 8-oxodG as a marker of oxidative damage in tracheoblasts, tracheae from L2 larvae were dissected and treated *ex vivo* with different concentrations of H_2_O_2_ (100μM, 500μM and 1mM) for 30 minutes and stained with an antibody against 8-oxodG. Robust staining was detected in trachea at 1mM H_2_O_2_ (Figure 4 A, n ≥ 6 tracheae per condition per experiment, n = 2) but no signal was detected in untreated tracheae. This showed that although 8-oxodG accumulation is responsive to oxidative stress, there is no 8-oxodG accumulation in G2-arrested tracheoblasts under normal conditions. An aspect of the 8-oxodG staining in tracheal cells is noteworthy and merits highlighting here. The accumulation of 8-oxodG in Tr2 tracheoblasts was cytoplasmic unlike the tracheal cells in other metameres, where 8-oxodG was observed in both cytoplasm and nucleus (Figure 4 B, n ≥ 6 tracheae per condition per experiment, n = 2). One reason for this difference could be that Tr2 DT are arrested in G2 and not engaged in DNA synthesis while cells in other metameres that are actively endocycling and replicating DNA.

**Figure 4:**
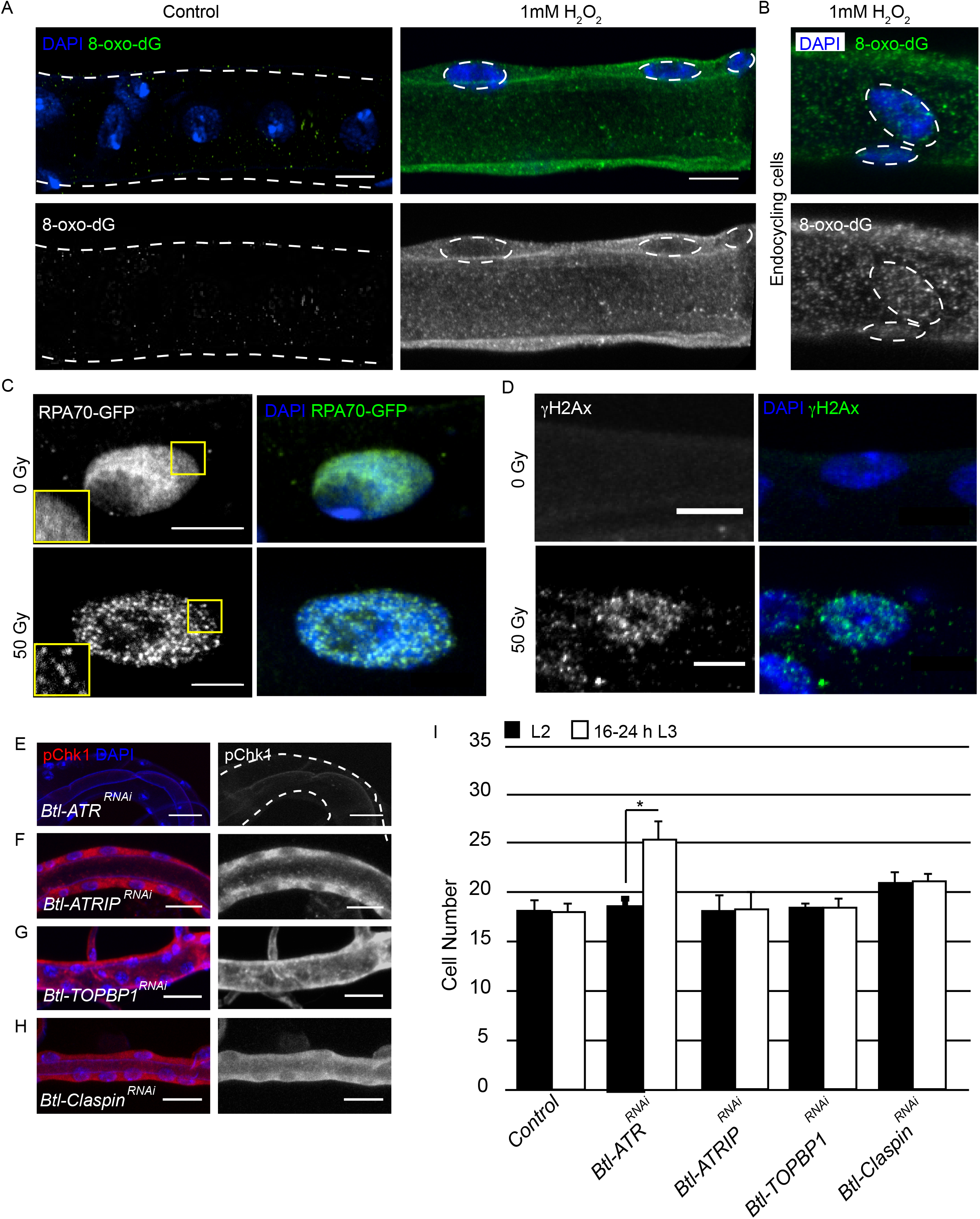
ATRIP, TOPBP1 and Claspin are not required for ROS mediated Chk1 activation in tracheoblasts. **(A-D**) Detailed analysis of DNA damage in Tr2 DT. Shown here are findings from three different reporters of genotoxic stress. **(A)** 8-Oxo-2’-deoxyguanosine (8-Oxo-dG) immunostaining in wild type *(btl-GAL4)* Tr2 DT in untreated tracheae (left panel) and tracheae exposed to 1mM H_2_O_2_ for 30 minutes *ex vivo* (right panel) at L2. **(B)** 8-Oxo-dG immunostaining in wild type *(btl-GAL4)* endocycling cells of the tracheae exposed to 1mM H_2_O_2_ for 30 minutes *ex vivo* at L2. **(C)** GFP immunostaining in larvae expressing RPA70-GFP. Shown in the figure are GFP immunostaining in non-irradiated larvae (top panel) and larvae exposed to 50 Gy of γ-radiation (bottom panel) at L2. **(D)** γ- H2AX^Ser139^ immunostaining in Tr2 DT in wild type *(btl-GAL4)* non-irradiated larvae (top panel) and larvae irradiated with 50 Gy of γ-radiation (bottom panel) at L2. **(E-H)** Analysis of the contribution of components of the DNA damage-dependent activation of ATR/Chk1 to Chk1 activation in Tr2 DT. Effects of the knockdown of *ATR, ATRIP, TOPBP1* and *Claspin* on pChk1 levels in Tr2 DT at L2. pChk1 immunostaining (red) in Tr2 DT in *Btl-ATR^RNAi^* (*btl-GAL4/UAS-ATR^RNAi^)* **(E)**, *Btl-ATRIP^RNAi^* (*btl-GAL4/UAS-ATRIP^RNAi^)* **(F)**, *Btl-TOPBP1^RNAi^* (*btl-GAL4/+; UAS-TOPBP1^RNAi^/+)* **(G)** and *Btl-Claspin^RNAi^* (*btl-GAL4/+; UAS-Claspin^RNAi^/+)* **(H)** larvae at L2. **(I)** Effects of knockdown of *ATR, ATRIP, TOPBP1* and *Claspin* on cell numbers in Tr2 DT. Graph shows numbers of Tr2 tracheoblasts in wild type *(btl-Gal4)*, *Btl-ATR^RNAi^* (*btl-GAL4/UAS-ATR^RNAi^)*, *Btl-ATRIP^RNAi^* (*btl-GAL4/UAS-ATRIP^RNAi^)*, *Btl-TOPBP1^RNAi^* (*btl-GAL4/+; UAS-TOPBP1^RNAi^/+)* and *Btl-Claspin^RNAi^* (*btl-GAL4/+; UAS-Claspin^RNAi^/+)* at L2 and 16-24 h L3 (mean values ± standard deviation, n ≥ 7 tracheae per condition per timepoint). Scale bars = 5 µm **(A-D)**, 10 µm (**E-H)**. Student’s paired t-test: *p<0.05.

Next we probed the incidence of single stranded DNA breaks in tracheoblats with the help of a strain that ubiquitously expresses RPA70-GFP (Blythe & Wieschaus, 2015). RPA 70 has been shown to be uniformly distributed in the nucleus under normal conditions and to form focal nuclear aggregates at sites of single strand DNA breaks (Blythe & Wieschaus, 2015). To validate RPA*70-GFP* as a marker for genotoxic stress in the tracheal system, L2 animals were exposed to either 0 (control) or 50 Gy of γ-irradiation and immunostained for GFP. Foci of GFP could be observed in the nuclei of tracheoblasts exposed to 50 Gy γ-irradiation (Figure 4 C, n ≥ 6 tracheae per condition per experiment, n = 3) but no foci were detected in untreated tracheae at the same stage. In a parallel set of experiments, we also examined levels of γ-H2AX in L2 animals under these conditions. We observed foci of nuclear γ-H2AX staining in tracheoblasts exposed to 50 Gy of γ-irradiation (Figure 4 D, n ≥ 6 tracheae per condition per experiment, n = 3) but not in untreated tracheae at the same stage. Taken together, our analysis of 8-oxodG, RPA70-GFP and γ- H2AX further showed that there is no detectable DNA damage in arrested tracheoblasts.

To probe the relationship between DNA damage and Chk1 activation in tracheoblasts, we also utilized a genetic approach. The mechanism for ATR/Chk1 activation in response to DNA damage has been characterized in some detail. These studies show that the activation of ATR/Chk1 requires three major proteins - ATR interacting protein (ATRIP, *mus-304*), topoisomerase II binding protein 1 (TOPBP1, *mus- 101*) and Claspin. Breaks in DNA that are bound by the single strand DNA binding protein RPA 70 recruits ATR via its partner ATRIP and, in turn, TOPBP1. This complex activates ATR and consequently, in a Claspin-dependent manner, Chk1 (Choi et al., 2010; Cimprich & Cortez, 2008; Delacroix et al., 2007; Lee et al., 2012; Xu & Leffak, 2010).

To confirm if ATRIP, TOPBP1 and Claspin are indeed required in tracheal cells for DNA damage-dependent phosphorylation of Chk1, we examined pChk1 levels post γ- irradiation in L2 tracheoblasts in which we simultaneous knocked down Duox and the aforementioned gene products. We exposed animals expressing *Btl-Duox^RNAi^,ATRIP^RNAi^, Btl-Duox^RNAi^,TopBP1^RNAi^* and *Btl-Duox^RNAi^, Btl-Duox^RNAi^,Claspin^RNAi^* to 50 Gy of γ-radiation and performed immunostaining for pChk1 1 h after irradiation (Figure 4-Figure supplement 1 A). pChk1 could be detected post irradiation in animals expressing *Btl-Duox^RNAi^* (Figure 5-Figure supplement1 C, n ≥ 6 tracheae per condition per experiment, n=3). We did not detect any pChk1 in *Btl-Duox^RNAi^,ATRIP^RNAi^*, *Btl-Duox^RNAi^,TOPBP1^RNAi^* and *Btl-Duox^RNAi^,Claspin^RNAi^* expressing animals at the same stages (Figure 4-Figure supplement 1 B-E, n ≥ 6 tracheae per condition per experiment, n = 2).This shows that DNA damage dependent phosphorylation of Chk1 in Tr2 DT cells requires ATRIP, TOPBP1 and Claspin.

To determine if any of the components of DNA damage dependent ATR activation are necessary for the phosphorylation of Chk1 in the trachea, we knocked down ATR, ATRIP, TOPBP1 and Claspin and probed the levels of pChk1 in L2 (Figure 4 E-H, n ≥6 tracheae per condition per experiment, n = 2). pChk1 staining of the tracheae from these animals revealed that the loss of ATR led to the loss of pChk1 (Figure 4 E). In contrast, the knockdown of ATRIP, TOPBP1 or Claspin did not lead to a loss of pChk1 in L2 (Figure 4 F-H). We also counted the number of cells in Tr2 DT at L2 and 16-24 h L3 in each of these genetic backgrounds. While the knockdown of ATR led to an increase in cell number at 16-24 h L3, knockdown of ATRIP, TOPBP1 and Claspin did not (Figure 4 I, n ≥ 7 tracheae per timepoint, Figure 4 – Source data 1). These data indicate that ATR-dependent activation of Chk1 in G2 arrested tracheoblasts cells does not require ATRIP, TOPBP1 and Claspin.

Collectively, these experiments reveal that Chk1 activation in tracheoblasts does not involve the DNA damage response pathway.

### H_2_O_2_ can rescue pChk1 levels in Duox deficient tracheoblasts

The next obvious question was to ask if Chk1 phosphorylation can be induced in Duox mutants by the addition of H_2_O_2._ To investigate this possibility, we examined levels of pChk1 in *Btl-Duox^RNAi^* tracheae after exposure to different concentrations of H_2_O_2_ for different periods of time. Tracheae from L2 animals were exposed to PBS or H_2_O_2_ (PBS) *ex vivo* and immunostained for pChk1 (Figure 5 A). We detected no pChk1 staining in tracheae exposed to buffer alone and robust staining in tracheae incubated with H_2_O_2_ (Figure 5 B-E, n ≥ 6 tracheae per condition per experiment, n=3). Interestingly, we noted that exposure to H_2_O_2_ for periods as short as 2 minutes was sufficient to restore levels of pChk1 in *Btl-Duox^RNAi^* tracheae (Figure5 E). We also probed pChk1 levels in *Btl-ATR^RNAi^* tracheae post H_2_O_2_ treatment and found that there was no pChk1 accumulation (Figure 5 G, n ≥ 6 tracheae per condition per experiment, n = 3). Together these data show that H_2_O_2_ treatment is sufficient to induce Chk1 phosphorylation in an ATR-dependent manner and that H_2_O_2_ can induce pChk1 in minutes.

**Figure 5:**
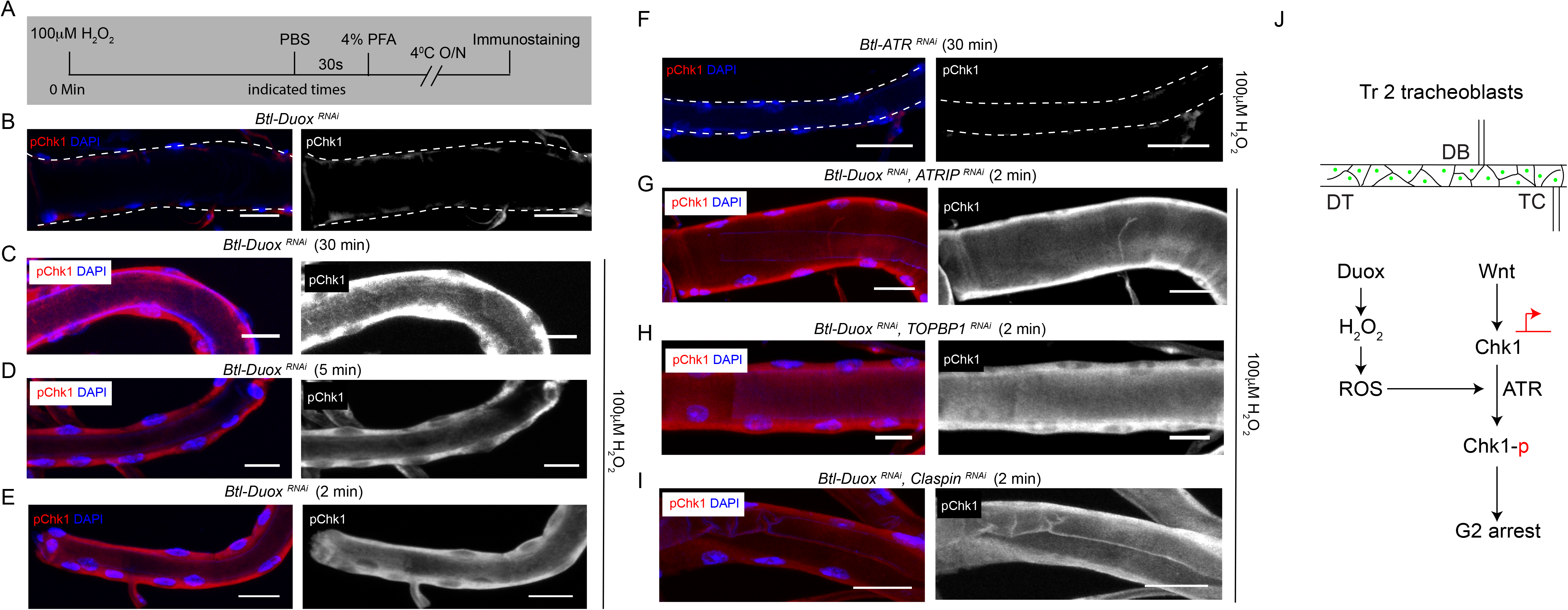
Incubation with H_2_O_2_ can restore pChk1 levels in Duox deficient tracheoblasts. **(A-E)** Kinetics of Chk1 phosphorylation upon exposure to H_2_O_2_ *ex vivo*. **(A)** Regimen for H_2_O_2_ treatment and analysis of pChk1 in Tr2 DT. pChk1 immunostaining (red) in Tr2 DT in *Btl-Duox^RNAi^ (btl-GAL4/+; UAS-Duox^RNAi^(32903)/+)* expressing tracheae- untreated **(B)** and treated with 100 µM H_2_O_2_ for 30 minutes **(C)**, 5 minutes **(D)** and 2 minutes **(E)** at L2. **(F)** Effect of knockdown of ATR on Chk1 activation in Tr2 DT upon exposure to H_2_O_2_ *ex vivo*. pChk1 immunostaining (red) in Tr2 DT in *Btl-ATR^RNAi^ (btl-GAL4/UAS-ATR^RNAi^)* tracheae treated with 100 µM H_2_O_2_ for 30 minutes. **(G-I** Effect of knockdown of *Duox* and *ATRIP*, *TOPBP1* or *Claspin* on pChk1 levels in Tr2 DT in tracheae exposed to 100 µM H_2_O_2_ at L2. pChk1 immunostaining (red) in Tr2 DT in *Btl-Duox^RNAi^, ATRIP^RNAi^* (*btl-GAL4/ UAS-ATRIP^RNAi^; UAS-Duox^RNAi^(32903)/+)* **(G)**, *Btl-Duox^RNAi^, TOPBP1^RNAi^ (btl-GAL4/+; UAS-Duox^RNA^(32903)^i^/UAS-TOPBP1^RNAi^)* **(H)** and *Btl-Duox^RNAi^, Claspin^RNAi^ (btl-GAL4/+; UAS- Duox^RNAi^(32903)/UAS-Claspin^RNAi^)* **(I)** tracheae treated with 100 µM H_2_O_2_ for 2 minutes at L2. **(J)** Model for the regulation of ATR/Chk1 activation in Tr2 DT. We propose that ATR, in the presence reactive oxygen species could activate Chk1 in a DNA damage independent manner leading to G2 arrest in Tr2 tracheoblasts. Scale bars = 10 µm

In an independent set of experiments, we examined the kinetics of DNA damage dependent activation of Chk1 in tracheoblasts. As described earlier, we exposed second instar larvae to 50Gy of γ-radiation and performed pChk1 immunostaining, at different timepoints post irradiation (Figure 5- Figure supplement 1 A,). We could detect pChk1 1 h hour post irradiation (Figure 5- Figure supplement 1 C, n ≥ 6 tracheae per condition per experiment, n=2) but not earlier (Figure 5- Figure supplement 1 D-E, n ≥ 6 tracheae per condition per experiment, n = 2). Here again, pChk1 induction in response to γ-radiation was dependent on ATR as pChk1 was not detected in tracheoblasts expressing *Btl-ATR^RNAi^* (Figure 5- Figure supplement 1 F, n ≥ 6 tracheae per condition per experiment, n = 2). These experiments suggest that the kinetics of Chk1 phosphorylation in response to H_2_O_2_ are significantly faster than in response to γ-radiation.

Next, we tested whether H_2_O_2_ could restore levels of pChk1 in the absence of ATRIP, TOPBP1 and Claspin in Btl-Duox*^RNAii^* expressing animals. To test this possibility, we exposed animals expressing *Btl-Duox^RNAi^,ATRIP^RNAi^*, *Btl-Duox^RNAi^,TOPBP1^RNAi^* and *Btl-Duox^RNAi^,Claspin^RNAi^* expressing tracheae to 100 μM H_2_O_2_. We performed Chk1 immunostaining 2 minutes after exposure. pChk1 immunostaining showed that the knockdown of ATRIP, TOPBP1 and Claspin did not prevent the activation of Chk1 (Figure 5 G-I, n ≥ 6 tracheae per condition per experiment, n = 2). This strongly suggests that the H_2_O_2_-dependent phosphorylation of Chk1 is independent on ATRIP, TOPBP1 and Claspin.

## DISCUSSION

ATR and Chk1 are essential for normal development in Drosophila and other animals. In this regard, both kinases are thought to serve as guardians of genomic integrity and loss of either is associated with increased genomic instability and catastrophic cell death. Our studies in the tracheal system reveal a different facet of ATR/Chk1 function during development. We have shown previously that the pathway is required to arrest tracheal progenitor cells in G2 and that the activation of ATR/Chk1 occurs in the absence of any detectable DNA damage. The findings presented here demonstrate that Duox-generated H_2_O_2_ is required for the activation of ATR/Chk1 axis in this context. We discuss below the possible mechanisms by which ROS can activate ATR/Chk1, the evidence that there are other mediators of non-canonical activation that are relevant to tracheal development, and the clinical implication of the findings.

The precedence for ROS-based activation of PIKK-family kinases like ATR, ATM and DNA-PK was set by studies on ATM. The formation of an inter-molecular disulfide bond between a cysteine residue located at the C-terminus of ATM was found to be essential for ATM homodimerization and activation. (Guo et al., 2010).The structure of Drosophila ATR is not known but the structure of human ATR in complex with ATRIP has been determined by cryoEM and the human ATR-ATRIP complex has been shown to dimerize (Rao et al., 2018). Using SWISS-MODEL and the kinase domain of human ATR as template, we modelled Drosophila ATR monomer (the sequence range modelled was 855-2517, which has 33.6 % sequence identity to human ATR). A model for the Drosophila ATR dimer was subsequently generated using the human ATR-ATRIP complex dimer as the template. This model shows that there are no exposed cysteines that are close enough to form inter-molecular disulfide bridges. Thus, there is a possibility that H_2_O_2_ regulates ATR in a different way than that has been proposed for ATM. Note that, it is not known whether ATR can also exist as a monomer sans ATRIP and thus the possibility that H_2_O_2_ can directly activate ATR by other mechanisms remains open. ROS can alter kinase activity either by modifying cysteine residues at or near the active site or more broadly, leading to conformational changes (Corcoran & Cotter, 2013a). These mechanisms may also be relevant here. ATR aside, ROS may also regulate Chk1 activation by modifying Chk1 in some manner or via recruitment of other factors. Future experiments will investigate these possibilities.

A comparison of the effects of ATR/Chk1 knockdown and SOD1 overexpression/Duox knockdown suggest that there are other (ROS-independent) mechanisms for the non-canonical activation of ATR/Chk1. We have shown that loss of either ATR or Chk1 leads to slow rate of cell division post mitotic reentry (Kizhedathu et al., 2018). Since there is no evidence for any DNA damage post mitotic re-entry and the knockdown of ATRIP, TOPBP1 and Claspin do not recapitulate the ATR/Chk1 mutant phenotype (Figure 4 – Figure supplement 2, n ≥ 7 tracheae per timepoint), the mechanism for the activation of ATR/Chk1 post mitotic re-entry is also non-canonical. Interestingly, neither the overexpression of SOD1 nor the knockdown of Duox recapitulate the mitotic defect observed in ATR/Chk1 mutants (Figure 1 J, Figure 2 F). This suggests that there are other non-canonical mediators of ATR/Chk1 activation that are relevant to the roles of these kinases during development.

The possibility that ROS can activate ATR/Chk1 without inducing DNA damage may be clinically relevant as well. It has been reported that ovarian cancer cells that express high levels of Duox1 and high levels of activated Chk1 are relatively Cisplatin-resistant (Meng et al., 2018). The authors suggest that high levels of Chk1 activation are critical for chemoresistance and that the high levels of Chk1 activation are likely the result of ROS-generated DNA damage. The findings presented here suggest that high ROS may independently activate Chk1 and contribute toward chemoresistance. Along the same lines, cancer cells that have high levels of ROS (Gu et al., 2018; Singh et al., 2020) and an active DNA damage repair pathway have been shown to be relatively radioresistant (Alsubhi et al., 2016; Wang et al., 2013). Here again, the enhanced radioresistance has been attributed to ROS-dependent DNA damage priming of the DNA damage response. We suggest that the ROS-based activation of ATR/Chk1 activation evidenced here may contribute toward the observed chemo/radioresistance of cancer cells.

## ACKNOWLEDGEMENTS

We thank Shigeo Hayashi, Eric F Wieschaus and M.J. Palladino for fly lines. The Central Imaging and Flow Cytometry Facility (CIFF) at inStem and Fly Facility at C-CAMP for their support. Support: Ramalingaswami Fellowship (Department of Biotechnology, Government of India, AG) and institutional funds from inStem (AK, PC).

## MATERIALS AND METHODS

### Key resources table

| Reagent type<br>(species) or resource | Designation | Source or<br>reference | Identifiers | Additional<br>information |
| --- | --- | --- | --- | --- |
| Genetic reagent<br>(*Drosophila melanogaster*) | *btl-GAL4* | (Shiga et al., 1996) | FLYB:<br>FBtp0001208 | This line was a gift from Dr. Shigeo Hayashi |
| Genetic reagent<br>(*Drosophila melanogaster*) | *UAS-Chk1<sup>RNAi</sup>* | VDRC | 110076 |  |
| Genetic reagent<br>(*Drosophila melanogaster*) | *RPA-70GFP* | (Blythe & Wieschaus, 2015) |  | This line was a gift from Dr. Eric F Wieschaus |
| Genetic reagent<br>(*Drosophila melanogaster*) | *UAS-Duox<sup>RNAi</sup>* | BDSC | 33975, 32903 |  |
| Antibody | Phospho-Chk1 (Ser345) | CST | 2348 | Rabbit monoclonal antibody (1:200) |
| Antibody | Anti-8-Hydroxy-2'-deoxyguanosine antibody | Abcam | ab48508 | Mouse monoclonal antibody (1:200) |
| Commercial assay or kit | Tyramide signal amplification system | Thermo Fisher Scientific | B40922 |  |

### Fly strains and Handling

The following strains were obtained from repositories: *UAS-Sod1* (24754), UAS*-Duox^RNAi^* (32903, 33975) UAS*-ATRIP^RNAi^* (61355), *UAS-TOPBP1^RNAi^* (43244), UAS*-Claspin^RNAi^* (32974) (Bloomington *Drosophila* Stock Center), *UAS-Chk1^RNAi^* (110076), *UAS-ATR^RNAi^* (103624) (Vienna *Drosophila* Resource Center). *UAS-Chk1* was generated in the in-house fly facility. The following strains were received as gifts: *btl-GAL4, UAS-MTS roGFP2*, *RPA70-GFP*. Strains were raised on a diet of cornmeal-agar and maintained at 25°C. All experiments were performed on animals raised at 25°C unless otherwise indicated.

### Larval Staging

Larval staging was performed as previously described (Guha & Kornberg, 2005) based on the morphology of the anterior spiracles. L2 larvae were collected and examined to identify animals that had undergone the L2-L3 molt in 8 hr intervals (0–8 h L3). 0–8 h L3 cohorts collected in this method were staged for subsequent time points.

### Immunostaining and Imaging

Animals were dissected in PBS and fixed for 30 min with 4% (w/v) Paraformaldehyde (PFA) in PBS. The following antisera were used for Immunohistochemical analysis: Chicken anti-GFP (Aves, 1:500), Rabbit anti-phospho Chk1 (CST, 1:200), Rabbit anti-pH3 (Millipore, 1:500), Mouse Anti-8-Hydroxy-2’-deoxyguanosine (Abcam, 1:200), and Alexa 488/568-conjugated Donkey anti-Chicken/Rabbit/Mouse secondary antibodies (Invitrogen, 1:200). Tyramide signal amplification was performed as per manufacturer recommendations for p-Chk1 detection. The following reagents were used as part of this protocol: Tyramide amplification buffer and Tyramide reagent (Thermofisher), Vectastain A and B and Biotinylated donkey anti Rabbit IgG (1:200, Vector Labs). Tracheal preparations were flat-mounted in ProLong Diamond Antifade Mountant with DAPI (Molecular Probes) and imaged on Zeiss LSM-780 laser-scanning confocal microscopes. Images were processed using ImageJ. For quantification of cell number, fixed specimens were mounted in ProLong Diamond Antifade Mountant with DAPI and the number of nuclei were counted from images collected with an Olympus BX 53 microscope. The DT of the second thoracic metamere was identified morphologically based on the cuticular banding pattern at anterior and posterior junctions.

### ROS Detection

Larvae of indicated stages were dissected in PBS, flipped inside out to expose the trachea, and incubated in 100 μM H_2_DCFDA (Thermofisher) for 30 minutes or 10 μM DHE (Thermofisher) for 5 minutes at room temperature. The larvae were then washed in PBS and fixed mildly in 4% PFA for 5 minutes. Tracheae were flat mounted in Prolong Diamond and imaged immediately. Mitochondrial ROS was measured using a genetically encoded redox-sensitive, mitochondrial-targeted GFP, MTS roGFP2 as described by (Liu et al., 2012). Briefly, animals expressing *Btl-MTS roGFP2* were dissected in PBS and the tracheae from these animals were flat mounted on a slide in PBS and imaged. Images were acquired at 535 nm emission using a 498-560 emission filter following excitation at 405 and 488 nm. The 405nm/488nm ratio was obtained using ImageJ software.

### RNA isolation and quantitative PCR

RNA extraction and qPCR were performed as described in (Kizhedathu et al., 2018). Primer sequences for *Chk1, Fz3, Wg, Wnt5, Wnt6, Wnt10, ATR, Duox* and *GAPDH* (internal control) are provided below. Relative mRNA levels were quantified using the formula RE = 2^− ΔΔCt^ method.

The following primer sets were used:

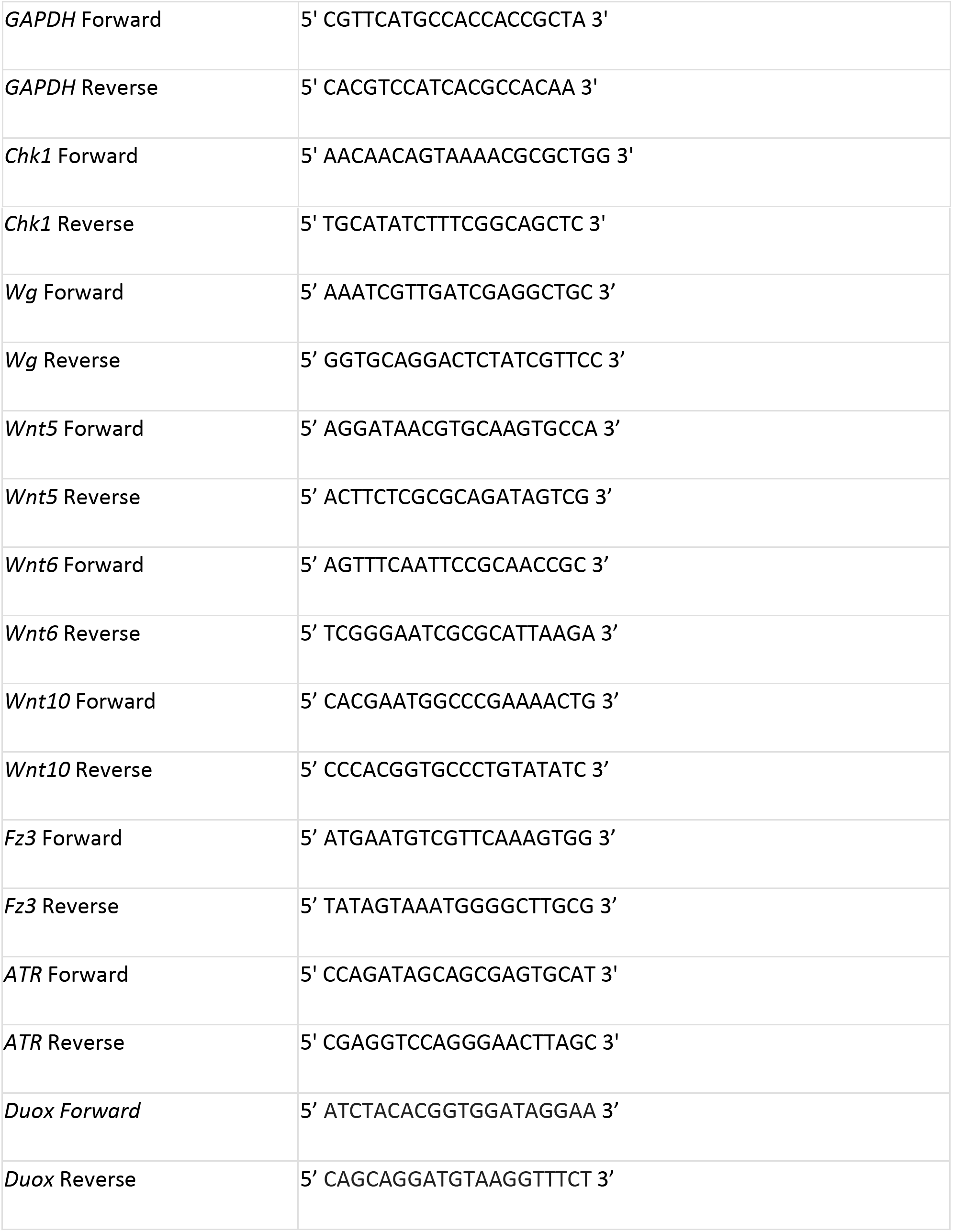

### γ-irradiation of larvae

Second instar larvae were exposed to 50Gy of γ-radiation at the rate of 2.56 Gy/minute using Blood Irradiator 2000 (Board of Radiation and Isotope Technology, DAE, Mumbai)). After irradiation, the larvae were transferred into media vials, maintained at 25°C for 3 h (detection of *RPA-70GFP* and γ−H2AX) or indicated time points (detection of pChk1) after which they were sacrificed.

### Hydrogen Peroxide treatment

Animals were dissected in PBS and flipped inside out to expose the tracheae. They were then incubated with specific concentrations of H_2_O_2_ in PBS at room temperature. For detection of 8-oxo-dG the larvae were immediately washed in PBS and fixed with 4% PFA and immunostaining was performed as described above. For detection of pChk1, the specimens were washed in PBS immediately and ice cold PFA was added. The samples were fixed overnight at 4°C. Immunostaining was then performed as indicated above.

## SUPPLEMENTARY FIGURE LEGENDS

**Figure 1 Supplement 1:**
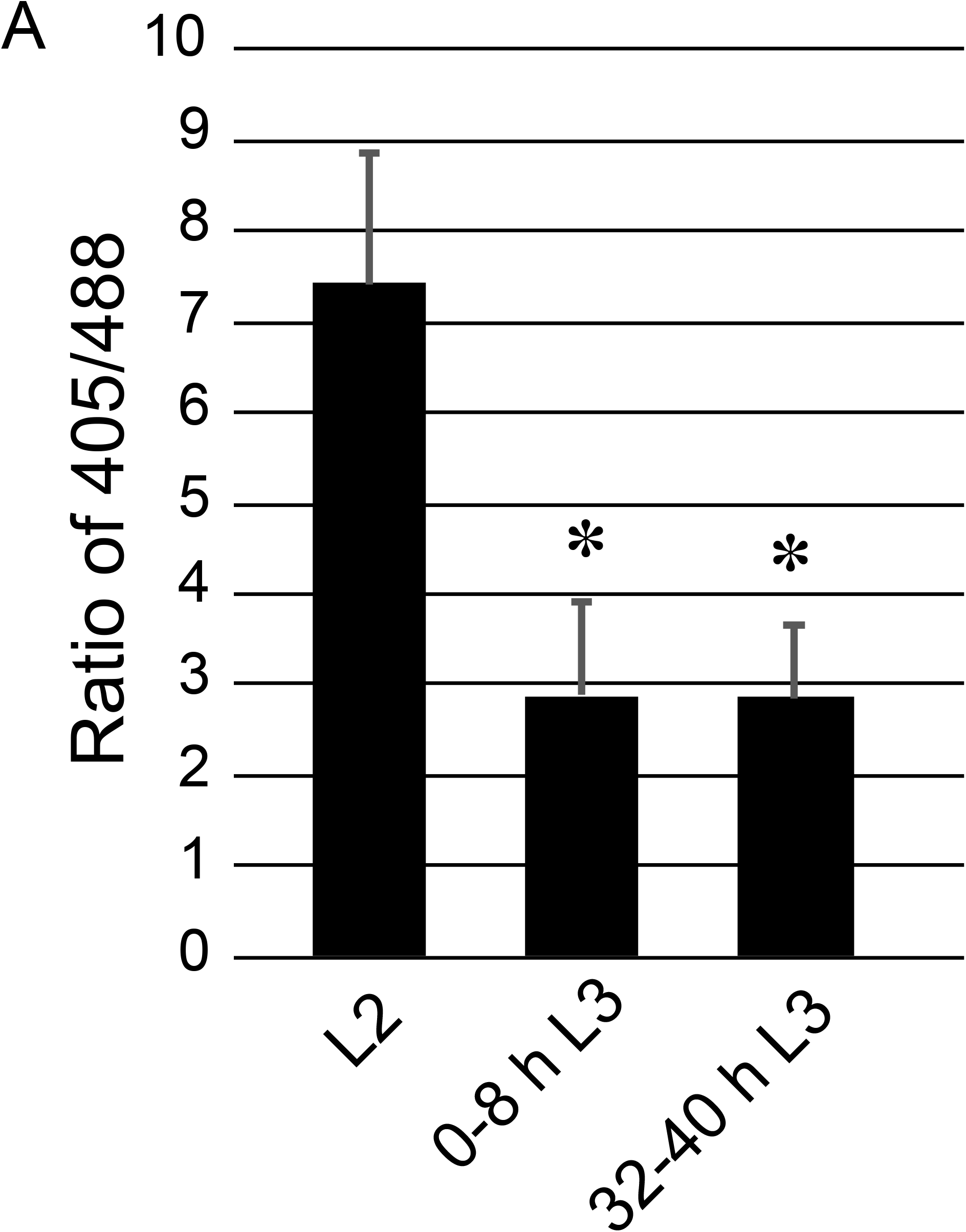
Quantification of ROS levels in the mitochondria. Graph shows ratio of fluorescence intensities of *Btl-MTS roGFP2* (*btl-GAL4/UAS-MTS roGFP2)* upon excitation at 405 and 488nm respectively as described by Liu et al., 2012 (also see methods). Fluorescence intensities were measured at L2, 0-8 h L3 and 32-40 h L3 (mean values ± standard deviation, n ≥ 6 tracheae per timepoint, Student’s paired t-test: *p<0.05).

**Figure 4 Supplement 1:**
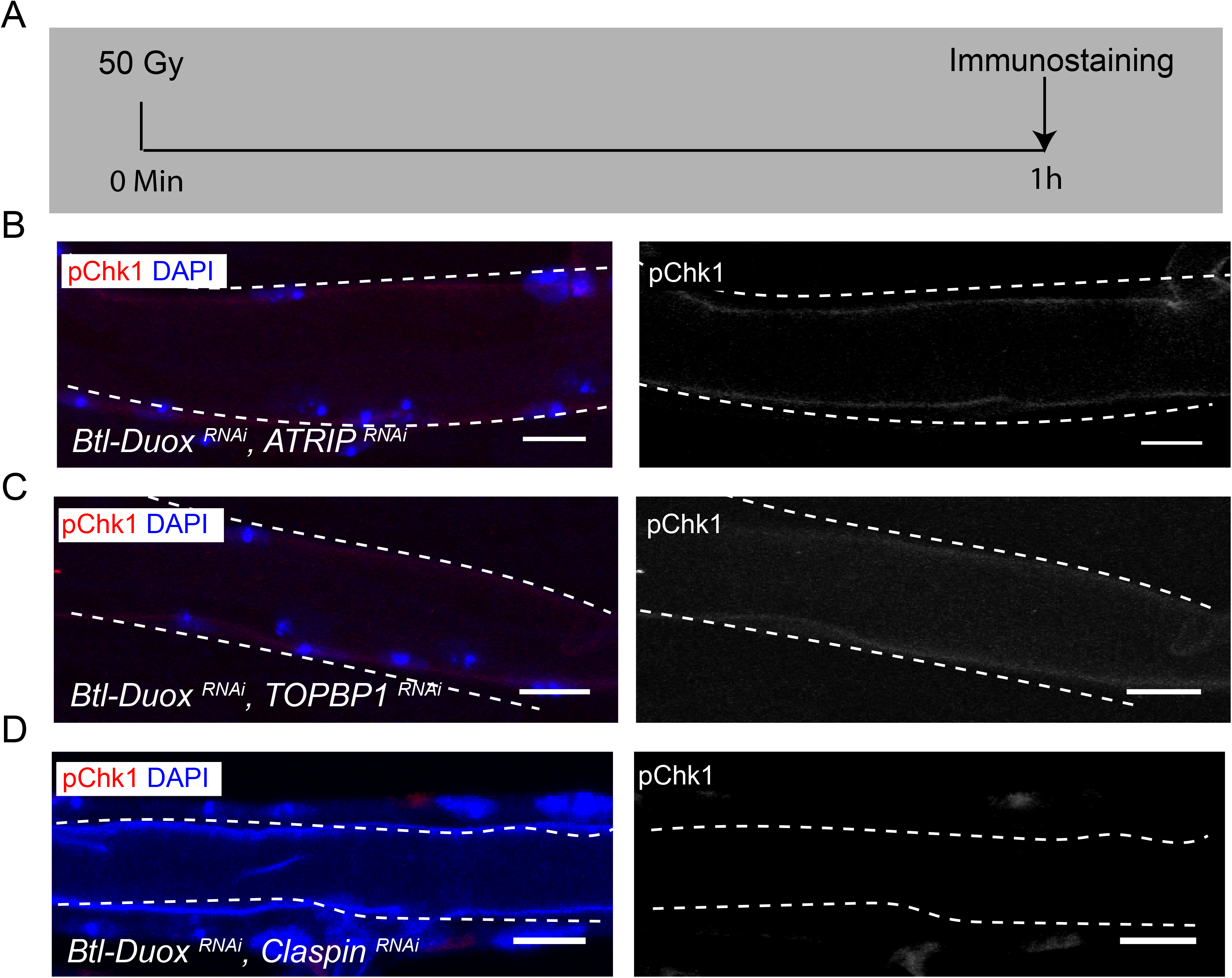
ATRIP, TOPBP1 and Claspin are required for DNA damage-dependent activation of ATR/Chk1. Effect of knockdown of *Duox* and *ATRIP*, *TOPBP1* or *Claspin* on pChk1 levels in Tr2 DT in larvae exposed to 50Gy of γ-radiation at L2. **(A)** Schematic describing the protocol for detection of pChk1 upon exposure to γ-radiation. **(B-D)** pChk1 immunostaining in Tr2 DT in **(B)** *Btl-Duox^RNAi^, ATRIP^RNAi^ (btl-GAL4/ UAS- ATRIP^RNAi^; UAS-Duox^RNAi^(32903)/+)*, **(C)** *Btl-Duox^RNAi^, TOPBP1^RNAi^ (btl-GAL4/+; UAS-Duox^RNAi^(32903)/UAS-TOPBP1^RNAi^)* and **(D)** *Btl-Duox^RNAi^, Claspin^RNAi^ (btl-GAL4/+; UAS- Duox^RNAi^(32903)/UAS-Claspin^RNAi^)* larvae at 1 h post exposure to 50 Gy of γ-radiation at L2. (n ≥ 6 tracheae per condition) Scale bars = 10 µm.

**Figure 4 Supplement 2:**
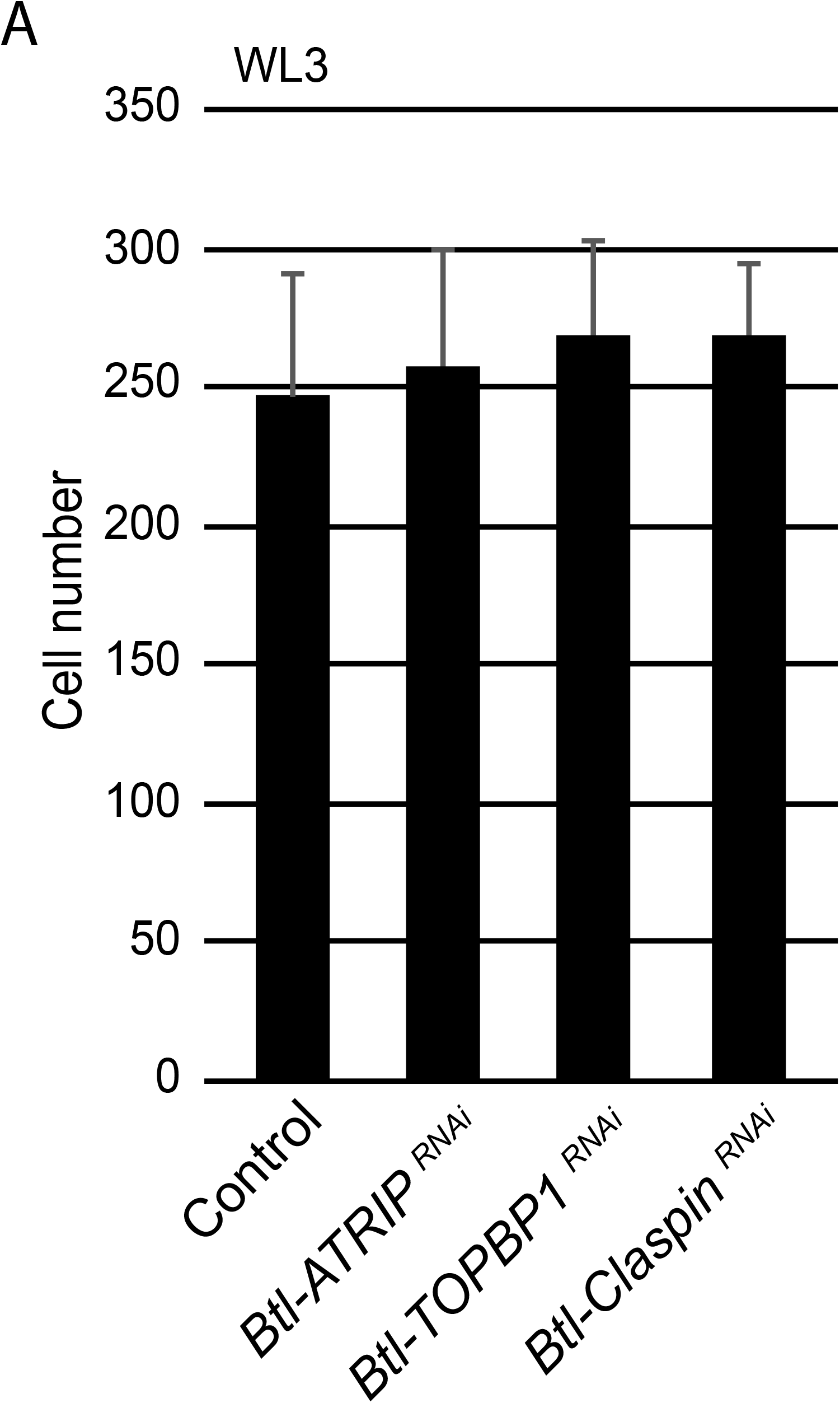
Loss of ATRIP, TOPBP1 and Claspin do not affect cell numbers at WL3. Effect of reduction of *ATRIP, TOPBP1* or *Claspin* on cell numbers in Tr2 DT at WL3. Graph shows cell numbers at WL3 in Control *(btl-GAL4), Btl-ATRIP^RNAi^* (*btl-GAL4/UAS- ATRIP^RNAi^)*, *Btl-TOPBP1^RNAi^ (btl-GAL4/+; UAS-TOPBP1^RNAi^/+)* and *Btl-Claspin^RNA^(btl-GAL4/+; UAS-Claspin^RNAi^/+)* (mean values ± standard deviation, n ≥ 7 tracheae).

**Figure 5 Supplement 1:**
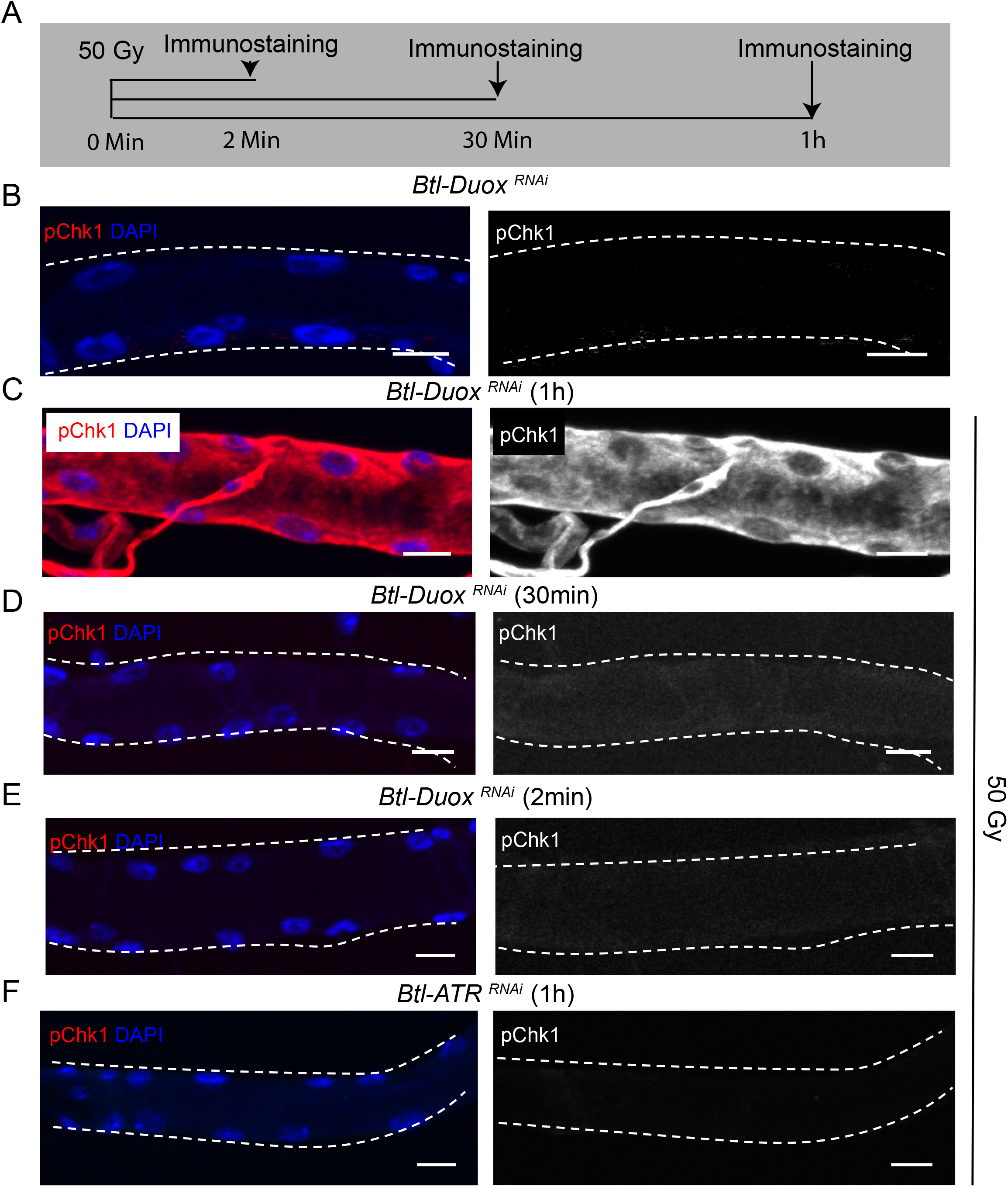
Exposure to γ-radiation can restore pChk1 levels in Duox deficient tracheoblasts. **(A-E)** Kinetics of Chk1 phosphorylation on exposure to γ-radiation. **(A)** Regimen for γ-irradiation and analysis of pChk1. pChk1 immunostaining (red) in Tr2 DT in *Btl-Duox^RNAi^ (btl-GAL4/+; UAS-Duox^RNAi^(32903/+)* in non-irradiated larvae **(B)** and larvae exposed to with 50 Gy of γ-radiation after 1 hour **(C)**, 30 minutes **(D)**, and 2 minutes **(E)** post irradiation at L2. **(F)** Effect of knockdown of ATR on Chk1 activation in Tr2 DT in larvae exposed to γ-radiation. pChk1 immunostaining (red) in Tr2 DT in *Btl-ATR^RNAi^ (btl- GAL4/UAS-ATR^RNAi^)* larvae exposed to 50 Gy of γ-radiation at 1 hour post exposure at L2. (n ≥ 6 tracheae per condition) Scale bars = 10 µm.

## Notes

### Competing Interest Statement

The authors have declared no competing interest.

